# A role for axon-glial interactions and Netrin-G1 signaling in the formation of low-threshold mechanoreceptor end organs

**DOI:** 10.1101/2022.06.16.496261

**Authors:** Shan Meltzer, Katelyn Comeau, Emmanuella Osei-Asante, Annie Handler, Qiyu Zhang, Chie Sano, Shigeyoshi Itohara, David D. Ginty

## Abstract

Low-threshold mechanoreceptors (LTMRs) and their cutaneous end organs convert light mechanical forces acting on the skin into electrical signals that propagate to the central nervous system. In mouse hairy skin, hair follicle-associated longitudinal lanceolate complexes, which are end organs comprising of LTMR axonal endings that intimately associate with terminal Schwann cell (TSC) processes, mediate LTMR responses to hair deflection and skin indentation. Here, we characterized developmental steps leading to the formation of Aβ RA-LTMR and Aδ-LTMR lanceolate complexes. During early postnatal development, Aβ RA-LTMRs and Aδ-LTMRs extend and prune cutaneous axonal branches in close association with nascent TSC processes. Netrin-G1 is expressed in these developing Aβ RA-LTMR and Aδ-LTMR lanceolate endings, and *Ntng1* ablation experiments indicate that Netrin-G1 functions in sensory neurons to promote lanceolate ending elaboration around hair follicles. The Netrin-G ligand (NGL-1), encoded by *Lrrc4c*, is expressed in TSCs, and ablation of *Lrrc4c* phenocopies the lanceolate complex deficits observed in *Ntng1* mutants. Moreover, NGL-1–Netrin-G1 signaling is a general mediator of LTMR end organ formation across diverse tissue types demonstrated by the fact that Aβ RA-LTMR endings associated with Meissner corpuscles and Pacinian corpuscles are also compromised in the *Ntng1* and *Lrrc4c* mutant mice. Thus, axon-glia interactions mediated by NGL-1–Netrin-G1 signaling promote LTMR end organ formation.

**Significance statement:** Our sense of touch is essential for fundamental tasks ranging from object recognition to social exchange. Yet, touch remains one of the least understood senses at the developmental level. Here, we investigate the formation of lanceolate complexes, which are mechanosensory end organs associated with hair follicles. The axons of touch sensory neurons innervating hairy skin extend into the skin at late embryonic and neonatal times, prune excessive branches during early postnatal development, and closely associate with non-myelinating glial cells, called terminal Schwann cells (TSCs) during formation of mature endings around hair follicles. Moreover, NGL-1 and its receptor Netrin-G1 mediate a molecular dialogue between nascent TSCs and sensory neuron axonal endings to promote mechanosensory end organ formation in both hairy and non-hairy (glabrous) skin.

## Introduction

Touch sensation is an integral component of our sensory experience, allowing us to perceive and respond to the physical world. Light touch is mediated by morphologically and physiologically distinct classes of low-threshold mechanosensory neurons (LTMRs), which detect a range of innocuous tactile stimuli and convey their signals from the skin to the central nervous system (Abraira and Ginty, 2013; Zimmerman et al., 2014; Jenkins and Lumpkin, 2017). The cell bodies of LTMRs are located in dorsal root ganglia (DRG) and cranial sensory ganglia. LTMRs have one axonal branch that extends to the skin and associates with different end organs and another branch that projects to the central nervous system and forms synapses onto second-order neurons in the spinal cord dorsal horn and brainstem (Fleming and Luo, 2013; Meltzer et al., 2021; Handler and Ginty, 2021). LTMRs have been classified as Aβ-, Aδ-, or C-LTMRs based on their action potential conduction velocities (Horch et al., 1977; Le Pichon and Chesler, 2014). Aβ-LTMRs are heavily myelinated and Aδ-LTMRs are lightly myelinated, exhibiting rapid and intermediate conduction velocities, respectively. C-LTMRs are unmyelinated and have a slow conduction velocity. LTMRs are also classified as slowly, intermediately, or rapidly adapting (SA-, IA-, and RA-, respectively) according to their firing patterns in response to sustained indentation of the skin (Handler and Ginty, 2021).

LTMR subtypes exhibit distinct intrinsic physiological properties and unique axonal endings associated with end organ structures across different skin types. The cutaneous axonal endings of Aβ RA-LTMR types include longitudinal lanceolate endings that enwrap hair follicles in hairy skin, terminate within Meissner corpuscles in glabrous skin, or form Pacinian corpuscles located in the deep dermis or around bones (Handler and Ginty, 2021). In mouse back hairy skin, Aβ RA-LTMR lanceolate endings form around guard hairs, which account for ∼1% of back skin hairs, and awl/auchene hairs. Aδ-LTMRs and C-LTMRs also form lanceolate endings, but unlike Aβ RA-LTMRs they are associated exclusively with awl/auchene and zigzag hairs (Li et al., 2011). These lanceolate ending structures are assumed to endow LTMRs with high sensitivity to hair deflection, skin indentation, and skin stroking (Handler and Ginty, 2021). Light and electron microscopic studies reveal that lanceolate endings are arranged parallel to the hair follicle’s long axis, and each lanceolate ending is encased by TSC processes (Kaidoh and Inoué, 2000; Li and Ginty, 2014; Takahashi-Iwanaga, 2000; Yamamoto, 1966). Similar to lanceolate endings in hairy skin, these endings are wrapped by non-myelinating Schwann cells called lamellar cells (Hashimoto, 1973; Idé, 1976; Neubarth et al., 2020; Pease and Quilliam, 1957; Spencer and Schaumburg, 1973; Takahashi-Iwanaga and Shimoda, 2003; Vega et al., 1994).

LTMR innervation of the skin occurs in parallel with skin and hair follicle morphogenesis (Fleming and Luo, 2013; Jenkins and Lumpkin, 2017; Meltzer et al., 2021; Olson et al., 2016). Beginning ∼E14.5, primary guard hair keratinocyte precursor cells elongate to form hair follicle placodes, and then proliferate and invaginate to form hair follicles (Paus et al., 1999). Secondary hair follicle development occurs in two waves: awl/auchene hairs develop at approximately E16.5, and zigzag hairs develop around birth (E18-P1) (Schneider et al., 2009). Developing hair follicles can release extrinsic cues to instruct the formation of lanceolate complexes. Keratinocytes on the caudal side of hair follicles express BDNF and control the polarized targeting of TrkB-expressing Aδ-LTMR endings to the caudal side of hair follicles (Rutlin et al., 2014). Moreover, hair follicle epidermal stem cells deposit EGFL6, an ECM protein, into the collar matrix, to regulate the proper patterning of lanceolate complexes (Cheng et al., 2018). However, the precise timing of LTMR innervation of hair follicles, whether LTMR axons undergo pruning during hair follicle innervation, the nature of the relationship between developing lanceolate endings and nascent TSCs, and molecular cues that instruct lanceolate ending morphological maturation remain unexplored.

Netrin-G1, encoded by *Ntng1*, is a member of the family of glycosyl-phosphatidylinositol (GPI)-anchored cell adhesion molecules. Netrin-G1 can promote synapse formation, microglial accumulation along axons, axonal outgrowth, and laminar organization of dendrites (Fujita et al., 2020; Lin et al., 2003; Matsukawa et al., 2014; Nishimura-Akiyoshi et al., 2007). Moreover, Netrin-G1 and its relative Netrin-G2 can localize to presynaptic membranes and instruct the specificity in synaptic connectivity (Matsukawa et al., 2014; Nishimura-Akiyoshi et al., 2007; Zhang et al., 2016a). In these contexts, Netrin-G1 is considered a receptor. The Netrin-G1 ligand, NGL-1, which is encoded by *Lrrc4c*, belongs to a family of postsynaptic adhesion molecules (Lin et al., 2003; Kim et al., 2006; Seiradake et al., 2011; Woo et al., 2009; Fujita et al., 2020). Mutations in both *Ntng1* and *Lrrc4c* have been implicated in neurological diseases, including Rett syndrome, schizophrenia, and autism (Aoki-Suzuki et al., 2005; Borg et al., 2005; Ohtsuki et al., 2008; Satterstrom et al., 2020). Yet, the functions of Netrin-G1 and NGL-1 in peripheral nervous system development have not been established.

Here, we used mouse genetic approaches to visualize Aβ RA-LTMRs and Aδ-LTMRs early during development (Luo et al., 2009; Rutlin et al., 2014), which allowed us to define the timing of hairy skin innervation and formation of their lanceolate complexes. Aβ RA-LTMR and Aδ-LTMR peripheral innervation patterns are established late embryonically and neonatally, exuberant axonal branches are pruned around birth, and newly formed lanceolate endings associate intimately with nascent TSCs. We show that Netrin-G1 signaling functions in somatosensory neurons to promote proper formation of Aβ RA-LTMR and Aδ-LTMR lanceolate complexes. *Lrrc4c*, encoding the Netrin-G1 ligand, NGL-1, is expressed in developing TSCs in hairy skin, and its deletion leads to similar, albeit milder, Aβ RA-LTMR and Aδ-LTMR lanceolate ending deficits. Moreover, we observed aberrant Meissner corpuscle and Pacinian corpuscle development in the absence of NGL-1–Netrin-G1 signaling. Our results delineate LTMR end organ developmental stages and reveal a role for NGL-1–Netrin-G1 signaling between TSCs and LTMR endings in mechanoreceptor end organ formation.

## Results

### Hair follicle innervation and pruning of Aβ RA-LTMR and Aδ-LTMR axons in hairy skin

We first investigated the timing of Aβ RA-LTMR and Aδ-LTMR innervation of hairy skin. For this, Aβ RA-LTMRs and Aδ-LTMRs were sparsely labeled by crossing Cre-dependent *Brn3a*^*f(AP)*^ reporter mice (Badea et al., 2009) with *Ret*^*CreER*^ or *TrkB*^*CreER*^ mice, respectively (Luo et al., 2009; Rutlin et al., 2014). To label developing Aβ RA-LTMR and Aδ-LTMR endings, pregnant females were treated with tamoxifen at E11.5 and E12.5, respectively. At E15.5, one day after guard hairs emerge, Aβ RA-LTMR axons were observed in the skin where they exhibited many small protrusions extending from the major branches, but they had not yet innervated nascent hair follicles (Figures 1A). Aβ RA-LTMR innervation of hair follicles was observed beginning ∼E17.5; these endings appeared as crescent-shaped axonal wrappings around nascent hair follicles. Many Aβ RA-LTMR axonal branches were observed at E17.5; most of these were pruned by postnatal day 3 (P3), a time point at which the number of branch points did not significantly differ from the number of innervated hair follicles (Figure 1B). In contrast, relative to Aβ RA-LTMRs, Aδ-LTMRs exhibited delayed innervation of hair follicles, as no Aδ-LTMR follicle innervation was observed until ∼P0 (Figures 1A and 1C). As with Aβ RA-LTMRs, Aδ-LTMRs extended extranumerary branches during their period of active hair follicle innervation at ∼P3, and branches that did not associate with hair follicles were pruned by P5 (Figure 1C). After P5, the number of branch points was still significantly more than the number of innervated hair follicles (Figure 1C), presumably because some hair follicles were innervated by more than one axonal branch of the same Aδ-LTMR neuron (Kuehn et al., 2019). Both Aβ RA-LTMRs and Aδ-LTMRs gradually increased the size of their innervation area during postnatal development as skin expansion occurred during growth of the pups (Figures 1D and 1E).

**Figure 1.**
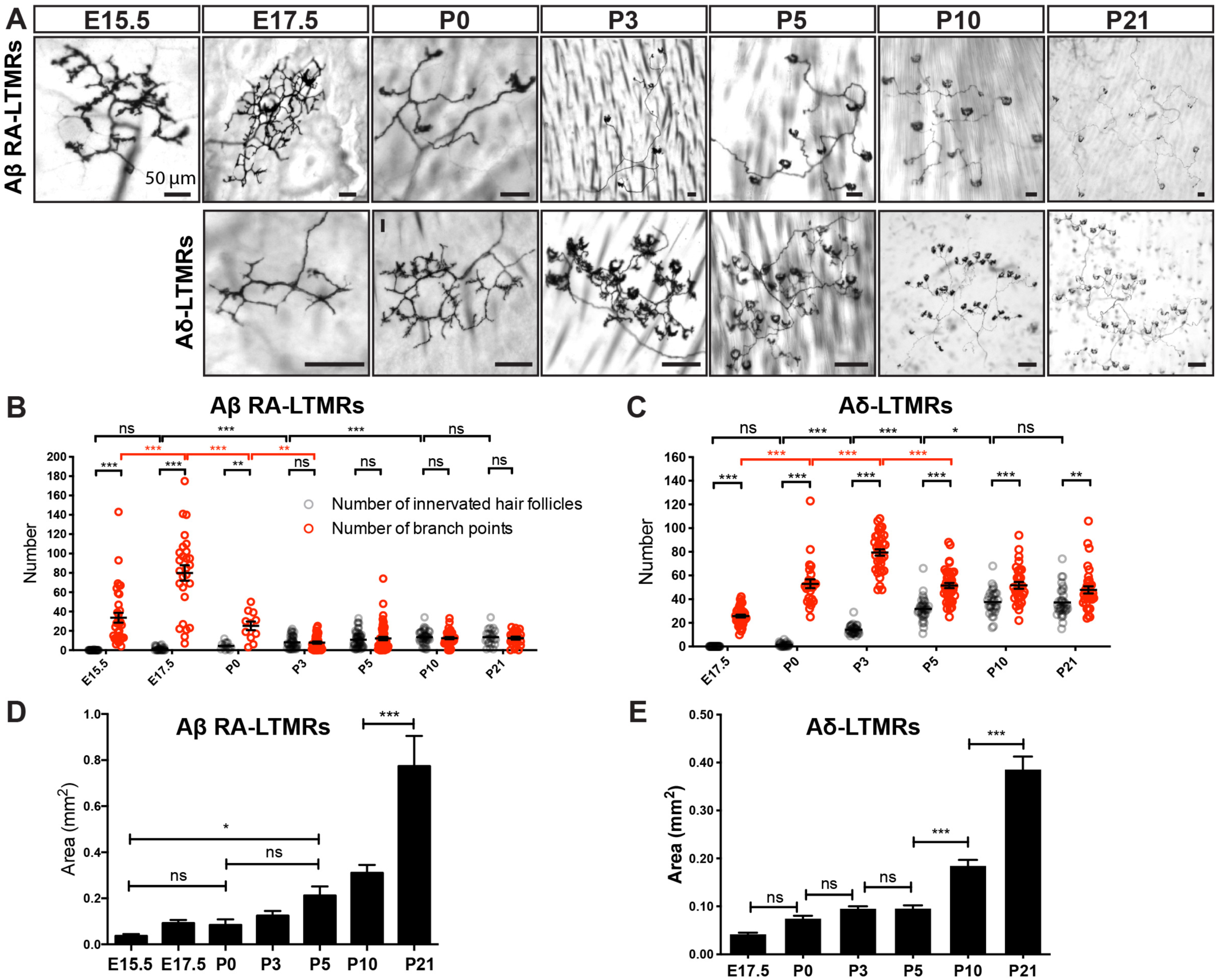
Peripheral arbors of Aβ RA-LTMRs and Aδ-LTMRs undergo pruning around birth. **(A)** Whole-mount AP staining of hairy skin to visualize peripheral terminals of individual Aβ RA-LTMRs and Aδ-LTMRs at different developmental time points and at P21. Scale bars represent 50 μm. **(B and C)** Quantification of the number of innervated hair follicles (grey) and branch points (red) of individual Aβ RA-LTMRs (B, n = 257 neurons from 37 animals) and Aδ-LTMRs (C, n = 210 neurons from 17 animals). Each dot represents a single neuron. Two-way ANOVA. **(D and E)** Quantification of the area of skin covered by individual Aβ RA-LTMRs (D) and Aδ-LTMRs (E). One-way ANOVA. ns, not significant, *p<0.05, **p<0.01, ***p<0.001

### Nascent Aβ RA-LTMR and Aδ-LTMR lanceolate endings closely interact with terminal Schwann cells during hair follicle innervation

We next examined the timing of Aβ RA-LTMR and Aδ-LTMR lanceolate complex formation. To visualize lanceolate endings around guard and non-guard hairs, we combined *Rosa26*^*LSL-tdTomato*^ (Ai14) with *Ret*^*CreER*^ and *TrkB*^*CreER*^ mice (Luo et al., 2009; Rutlin et al., 2014) and performed wholemount immunostaining at different time points during postnatal development (Figure 2A). At P0, nascent Aβ RA-LTMR lanceolate endings were observed around guard and non-guard hair follicles, while Aδ-LTMRs only formed crescent endings around non-guard hair follicles and did not yet exhibit lanceolate endings (Figures 2A and 2B). Lanceolate endings around guard hairs were already significantly longer than those around non-guard hairs, which is consistent with the earlier formation and maturation of guard hairs (Figure 2B). By P5, lanceolate endings from both Aβ RA-LTMRs and Aδ-LTMRs were observed, and lanceolate endings around guard hairs were significantly longer than those around non-guard hairs (Figure 2B). By P10, the length of Aβ RA-LTMR lanceolate endings around non-guard hairs was comparable to those of Aδ-LTMRs. Lanceolate endings continued to extend during postnatal development and achieved mature morphology by P21 (Figures 2A and 2B).

**Figure 2.**
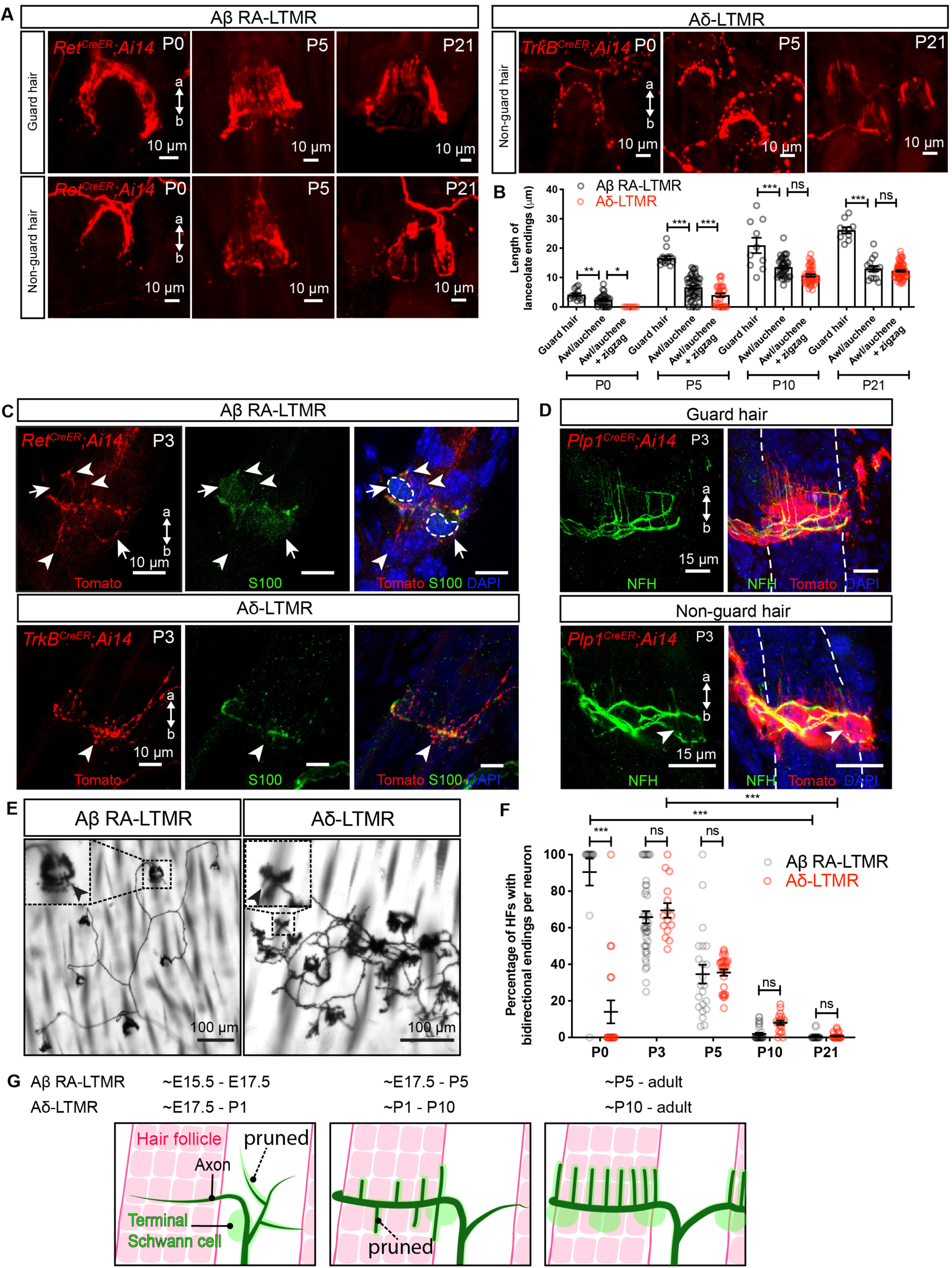
Lanceolate endings of Aβ RA-LTMRs and Aδ-LTMRs emerge around hair follicles neonatally and closely associate with terminal Schwann cells. **(A)** Whole-mount immunostaining images of back hairy skin showing Aβ RA-LTMRs in *Ret*^*CreER*^*;Ai14* animals and Aδ-LTMRs in *TrkB*^*CreER*^*;Ai14* animals at P0, P5 and P21. Guard hairs were identified by TROMA-I (Krt8) staining that labels Merkel cells that assemble into touch domes. **(B)** Quantification of the length of lanceolate endings associated with hair follicles. Aβ RA-LTMRs (black) and Aδ-LTMRs (red) at P0, P5, P10 and P21, showing that Aβ RA-LTMR lanceolate endings innervating guard hairs develop earlier than the lanceolate endings innervating awl/auchene and zigzag hairs (together grouped as non-guard hairs). At least 3 animals were used for each time point. Each dot represents lanceolate ending length around a hair follicle. One-way ANOVA was used for each time point. **(C)** Back hairy skin sections from *Ret*^*CreER*^*;Ai14* animals (n = 3 animals) and *TrkB*^*CreER*^*;Ai14* animals (n = 3 animals) were stained with an anti-Tomato antibody to label Aβ RA-LTMRs and Aδ-LTMR axonal terminals around hair follicles at P3. S100 (green) staining labels TSCs and DAPI labels nuclei. White arrowheads point to some of the nascent lanceolate endings growing towards the apical and basal side of the skin. White arrows and dotted lines denote cell bodies of TSCs. **(D)** Whole-mount immunostaining images from *Plp1*^*CreER*^*;Ai14* animals (n = 3 animals), which were stained with an anti-Tomato antibody to label TSCs at P3. NFH (green) labels lanceolate endings of Aβ RA-LTMRs. White arrowhead points to nascent lanceolate endings growing towards the basal side of the skin. **(E)** Whole-mount alkaline phosphatase staining of hairy skin reveals peripheral terminals from individual Aβ RA-LTMRs and Aδ-LTMRs sparsely labeled at P3. Insets showing a high magnification region of lanceolate endings around hair follicles. Black arrowheads point to nascent lanceolate endings extending towards the basal side of the skin. **(F)** Quantification of the percentage of hair follicles with bidirectional endings per neuron for Aβ RA-LTMRs and Aδ-LTMRs at P0, P5, P10 and P21. Each dot represents the percentage of hair follicles with bidirectional endings measured for a single neuron. At least 3 animals were used for each time point. Two-way ANOVA was used. **(G)** Summary of peripheral innervation steps for Aβ RA-LTMRs and Aδ-LTMRs. Black represents sensory axons, green cells represent terminal Schwann cells, and hair follicles are depicted in red. Dotted lines show that some sensory neuron branches and lanceolate endings are pruned during development. a, apical. b, basal. ns, not significant, *p<0.05, **p<0.01, ***p<0.001

Mature LTMR lanceolate endings are closely associated with TSCs, whose cell bodies reside at the base of lanceolate complexes (Halata, 1993; Li and Ginty, 2014). To visualize the temporal and morphological relationships between lanceolate axonal endings and TSCs during lanceolate complex formation, wholemount immunostaining with S100, a peripheral glia marker, and tdTomato was performed using skin from P3 *Ret*^*CreER*^; *Rosa26*^*LSL-tdTomato*^ and *TrkB*^*CreER*^; *Rosa26*^*LSL-tdTomato*^ animals. Although S100 immunolabeling was faint at P3, some Aβ RA-LTMR and Aδ-LTMR lanceolate endings were closely associated with this glial marker at this time point (Figure 2C). As a complementary approach, *Plp1*^*CreER*^; *Rosa26*^*LSL-tdTomato*^ animals were generated to genetically label TSCs and visualize their processes. Similar to lanceolate axons, TSCs exhibited exuberant processes around hair follicles at P3. Neurofilament H (NFH)-positive Aβ RA-LTMR lanceolate endings associated with both guard and non-guard hairs were closely associated with TSC processes at this age (Figure 2D), with 97.9% of NFH^+^ lanceolate endings around guard hairs enwrapped by TSC protrusions (140 lanceolate endings from 3 animals). Thus, lanceolate axonal endings are intimately associated with TSC processes during the period of lanceolate complex formation.

### Aβ RA-LTMRs and Aδ-LTMRs prune basal-orienting lanceolate endings during early postnatal development

Mature lanceolate endings orient exclusively towards the apical side of the epidermis (Figure 2A) (Halata, 1993; Li and Ginty, 2014; Rutlin et al., 2014). Interestingly, we frequently observed lanceolate endings extending towards the basal side of the epidermis in P3 skin (Figures 2C and 2D), suggesting that these basal-orienting lanceolate endings are pruned later during postnatal development. To assess the prevalence and elimination of these basal-orienting lanceolate endings, we examined sparsely labeled Aβ RA-LTMRs and Aδ-LTMRs in hairy skin using wholemount alkaline phosphatase labeling and high magnification visualization. We quantified the percentage of hair follicles with lanceolate endings pointing to both apical and basal side of the epidermis (bidirectional lanceolate endings) for individual neurons across postnatal development (Figures 2E and 2F). At P21, lanceolate endings wrapping around hair follicles pointed exclusively towards the apical side of the epidermis (Figure 1A). Conversely, at P0, most hair follicles innervated by Aβ RA-LTMRs exhibited lanceolate endings with both apical and basal orientations. Moreover, although at P0 most Aδ-LTMRs had just begun to extend lanceolate endings, those that did often displayed bidirectional orientation (Figure 2F). The percentages of hair follicles with bidirectional endings gradually decreased for both Aβ RA-LTMRs and Aδ-LTMRs during subsequent postnatal development and bidirectional endings were eventually gone by P21 (Figure 2F). The temporal similarity in the pruning of basal-orienting endings across these two lanceolate ending neurons suggests that there may be a common cue regulating the elimination of basal-orienting lanceolate endings.

Thus, despite a difference in their timing of skin innervation, Aβ RA-LTMRs and Aδ-LTMRs share developmental features (Figure 2G). First, both populations exhibit excessive branching and prune axonal branches that do not innervate hair follicles, with Aβ RA-LTMRs undergo pruning earlier than Aδ-LTMRs. Second, both LTMRs actively extend axons to innervate hair follicles while greatly expanding their peripheral arbor area during growth of the body. Third, both have nascent lanceolate protrusions that associate intimately with TSC processes. Fourth, both exhibit apical- and basal-oriented lanceolate extensions and eliminate the basal extensions to achieve their mature morphological properties by P21.

### Netrin-G1 is expressed in developing and mature Aβ RA-LTMRs and Aδ-LTMRs

To begin to test the hypothesis that common molecular mechanisms may function across LTMR types during the assembly of lanceolate complexes, we searched published datasets for cell-adhesion molecules and axon guidance proteins expressed in developing Aβ RA-LTMRs and Aδ-LTMRs, but not in nociceptors or proprioceptors which do not form lanceolate complexes (Sharma et al., 2020; Zheng et al., 2019). Netrin-G1 stood out in this analysis because it is expressed in both Aβ RA-LTMRs and Aδ-LTMRs throughout development and in adults (Sharma et al., 2020; Zheng et al., 2019). Netrin-G1 plays critical roles in many aspects of neural development, including functioning as a synaptic cell adhesion molecule to promote excitatory synapse formation, regulating formation of dendritic laminar structures, and axonal outgrowth (Kim et al., 2006; Lin et al., 2003; Matsukawa et al., 2014; Nishimura-Akiyoshi et al., 2007). Thus, we hypothesized that Netrin-G1 may orchestrate LTMR lanceolate ending complex formation or other features of Aβ RA-LTMR and Aδ-LTMR axonal development.

We first verified expression of Netrin-G1 in P40 LTMRs using a Netrin-G1 antibody (Vue et al., 2013). All Netrin-G1 positive DRG cells were large diameter and NFH^+^, consistent with their expression in NFH^+^ Aβ RA-LTMRs and Aδ-LTMRs (Figure 3A). As a control for Netrin-G1 antibody specificity, we used *Ntng1*^*-/-*^ mutants and found no antibody signal (Figures 3A-3D). Netrin-G1 protein was also detected in lanceolate endings around hair follicles as well as Aβ RA-LTMR axon terminals in Meissner corpuscles of glabrous skin (Figures 3B-3C). Netrin-G1 signal was also detected in Aβ field-LTMR circumferential endings in hairy skin and at a low level in lamellar cells associated with Meissner corpuscles (Figures 3B-3C). In the spinal cord, Netrin-G1 was detected in the region below lamina II (marked by IB4), consistent with the central projection pattern of Aβ RA-LTMRs and Aδ-LTMRs (Figure 3D). To ask whether Netrin-G1 is expressed in developing Aβ RA-LTMRs and Aδ-LTMRs, we performed Netrin-G1 immunostaining with DRGs and skin from *Ret*^*CreER*^; *Ai14* and *TrkB*^*CreER*^; *Ai14* animals. Indeed, Netrin-G1 was present in both the cell bodies and lanceolate endings of genetically labeled Aβ RA-LTMRs and Aδ-LTMRs at P3 (Figures 3E and 3F). Thus, Netrin-G1 is expressed in developing Aβ RA-LTMRs and Aδ-LTMRs and localizes to their axonal endings during the period of lanceolate ending maturation.

**Figure 3.**
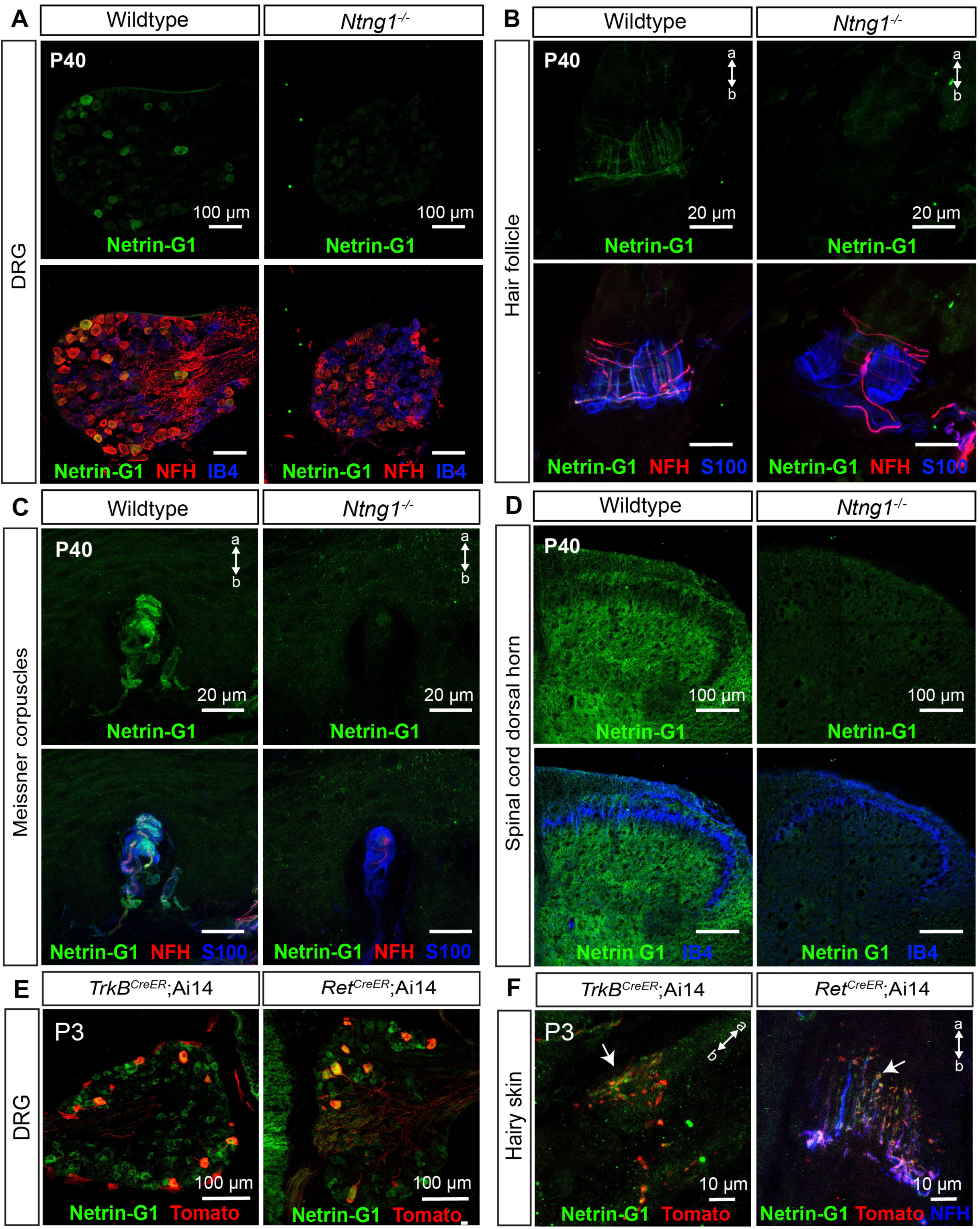
Netrin-G1 is expressed in developing and adult Aβ RA-LTMRs and Aδ-LTMRs. **(A-D)** IHC images of DRG sections from control and *Ntng1*^*-/-*^ animals (n = 3 animals per genotype) at P40, showing expression of Netrin-G1 in NFH^+^ DRG neurons (A), lanceolate and circumferential endings around hair follicles (B), Aβ RA-LTMRs innervating Meissner corpuscles (C), and spinal cord (D). Netrin-G1 signal (green) is absent in *Ntng1*^*-/-*^ animals. IB4 labels a large subset of non-peptidergic sensory neurons and S100 labels TSCs. **(E and F)** Thoracic DRG and back hairy skin sections obtained from P3 *Ret*^*CreER*^*;Ai14* animals (n = 3 animals) and *TrkB*^*CreER*^*;Ai14* animals (n = 3 animals) were stained with Netrin-G1 and Tomato to visualize Aβ RA-LTMRs and Aδ-LTMRs. NFH (blue) labels lanceolate endings of Aβ RA-LTMRs. a, apical. b, basal.

### Netrin-G1 regulates lanceolate ending formation in Aβ RA-LTMRs and Aδ-LTMRs

We next tested the hypothesis that Netrin-G1 contributes to Aβ RA-LTMR skin innervation and lanceolate complex formation using *Ntng1* knockout mice. For this, lanceolate endings around guard hairs of P40 *Ntng1*^*-/-*^, *Ntng1*^*+/-*^, and littermate control mice were visualized with NFH and Tuj1 immunostaining (Figures 4A and S1). The number of lanceolate endings around guard hairs in *Ntng1*^*-/-*^ mice was reduced compared to controls (Figure 4B and Figure S1). A comparable number of NFH^+^ neurons was observed in DRGs of *Ntng1*^*-/-*^ and control mice, suggesting that the reduction of lanceolate endings was not caused by loss of DRG neurons (Figure S2). Moreover, lanceolate endings in *Ntng1*^*-/-*^ mice showed a substantial number of enlargements in their distal tip regions (Figures 4A and 4C). Electron microscopy micrographs show that these enlarged endings had reduced TSC wrappings and disorganized intracellular structures (i.e. more vesicles) (Figure 4D). These enlarged endings with aberrant ultrastructural features indicate that the structural integrity of lanceolate endings is compromised in *Ntng1*^*-/-*^ mice. In addition, *Ntng1*^*+/-*^ mice exhibited an intermediate phenotype in both the number of lanceolate endings and the number of enlarged endings around guard hairs, suggesting that a precise level of *Ntng1* expression is critical for lanceolate ending formation (Figures 4B, 4C and S1). To determine if similar deficits exist in Aδ-LTMRs, we visualized Aδ-LTMR lanceolate endings using *TrkB*^*GFP*^ mice (Rutlin et al., 2014). As in Aβ RA-LTMRs, the number of Aδ-LTMR lanceolate endings was decreased and the number of lanceolate ending enlargements around non-guard hairs was increased in *Ntng1*^*-/-*^ mice, indicating that Netrin-G1 regulates lanceolate complex formation in both Aβ RA-LTMRs and Aδ-LTMRs (Figures 4E-4G).

**Figure 4.**
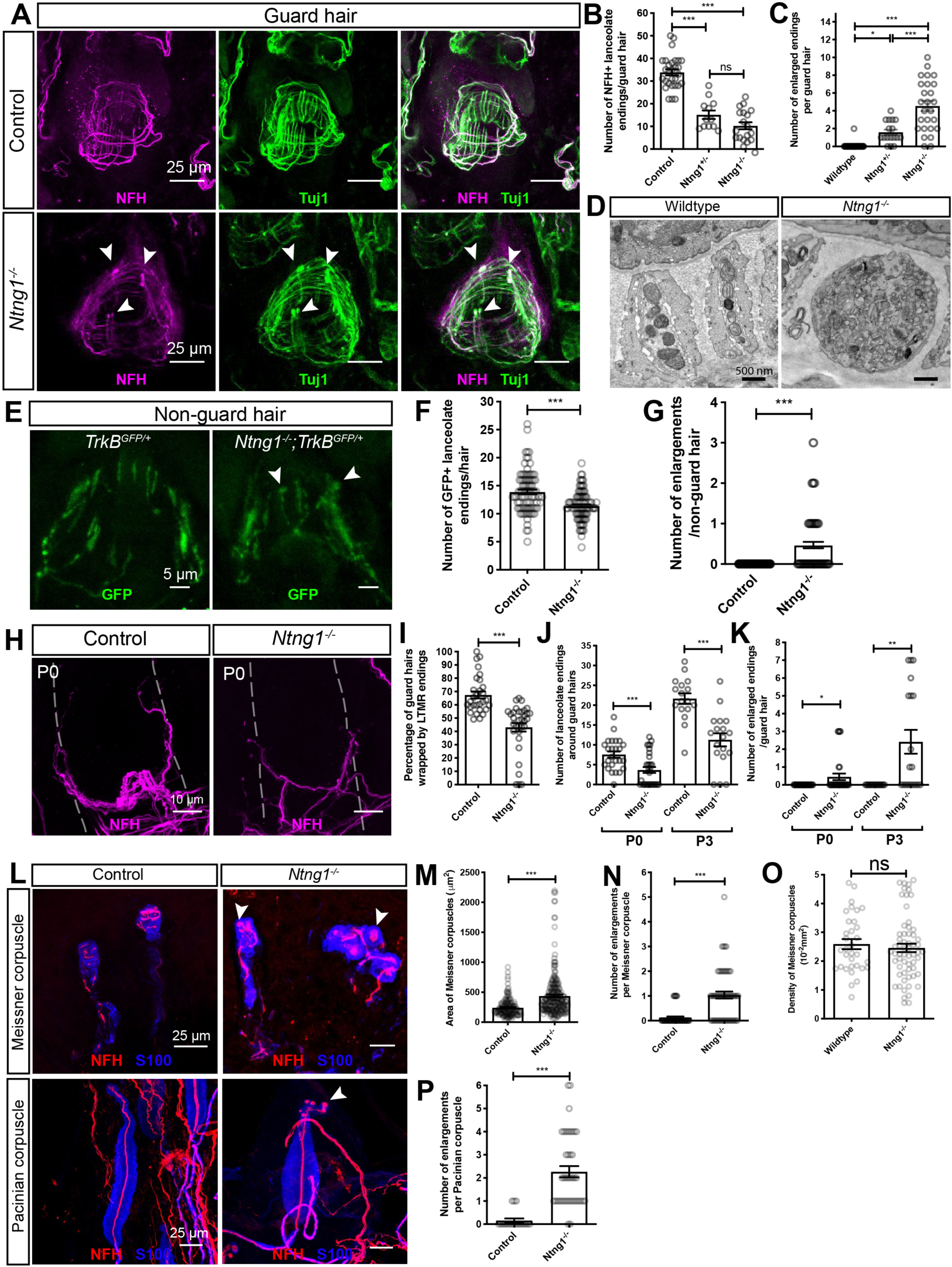
Netrin-G1 regulates the formation of lanceolate endings, Meissner corpuscles and Pacinian corpuscles. **(A)** Whole-mount immunostaining images of guard hairs in back hairy skin of adult control and *Ntng1*^*-/-*^ animals. Aβ RA-LTMRs lanceolate endings are marked by NFH (magenta) and Tuj1 (green) labeling. **(B and C)** Quantification of the number of NFH^+^ lanceolate endings (B) and the number of enlarged endings (C) per guard hair for Aβ RA-LTMRs in control (n = 7 animals), *Ntng1*^*+/-*^ (n = 2 animals) and *Ntng1*^*-/-*^ (n = 3 animals) animals, showing fewer Aβ RA-LTMR lanceolate endings in *Ntng1*^*+/-*^ and *Ntng1*^*-/-*^ animals. Each dot represents a single hair follicle. One-way ANOVA test. **(D)** A transmission electron microscopic image of a cross section through a lanceolate complex of a guard hair follicle for wildtype and *Ntng1*^*-/-*^ animals. **(E)** Whole-mount immunostaining images of non-guard hairs in back hairy skin in *TrkB*^*GFP/+*^ and *Ntng1*^*-/-*^; *TrkB*^*GFP/+*^ animals. Aδ-LTMRs lanceolate endings are marked by GFP labeling. **(F and G)** Quantification of the number of GFP^+^ lanceolate endings (F) and the number of enlarged endings (G) in *TrkB*^*GFP/+*^ (n = 3 animals) and *Ntng1*^*-/-*^; *TrkB*^*GFP/+*^ (n = 3 animals) animals, showing similar deficits in lanceolate ending formation with non-guard hairs. Student’s unpaired t test. **(H)** Whole-mount immunostaining images of guard hairs in back hairy skin of P0 control and *Ntng1*^*-/-*^ animals. **(I)** Quantification of the percentages of guard hairs wrapped by NFH^+^ lanceolate endings at P0 in control and *Ntng1*^*-/-*^ animals. Student’s unpaired t test. **(J and K)** Quantification of the number of NFH^+^ lanceolate endings (J) and the number of enlarged endings (K) per guard hair for Aβ RA-LTMRs in control (n = 3 animals for P0, n = 5 animals for P3) and *Ntng1*^*-/-*^ (n = 3 animals for P0, n = 5 animals for P3) animals. Student’s unpaired t test. **(L)** Representative IHC images of forepaw glabrous skin sections and Pacinian corpuscles. Meissner corpuscles and Pacinian corpuscles are labeled by S100 (blue) for visualizing lamellar cells and NFH (red) for visualizing Aβ RA-LTMRs. Arrowheads point to axonal enlargements. **(M-O)** Quantification of the area (M), number of enlargements (N), and density (O) of Meissner corpuscles in the epidermis of control (n = 4 animals) and *Ntng1*^*-/-*^ (n = 4 animals) mice. Each dot represents a single skin section (O). Student’s unpaired t test. **(P)** Quantification of the number of enlargements per Pacinian corpuscle from control (n = 3 animals) and *Ntng1*^*-/-*^ (n = 3 animals) mice. Student’s unpaired t test. Each dot represents a single Pacinian corpuscle. ns, not significant, *p<0.05, **p<0.01, ***p<0.001

To determine whether the paucity of lanceolate complexes in *Ntng1*^*-/-*^ adult mice reflects a developmental alteration in lanceolate ending formation or a loss of mature lanceolate endings in adults, we next assessed the timing of hair follicle innervation deficits during postnatal development in the mutants. At P0, *Ntng1*^*-/-*^ mice exhibited a significant reduction in the percentage of guard hair follicles wrapped by Aβ RA-LTMRs (Figures 4H and 4I). While control animals start to exhibit nascent lanceolate endings around guard hairs at P0 and continue at P3, lanceolate ending extension in *Ntng1*^*-/-*^ mutants was significantly reduced at these times (Figure 4J). Similarly, enlarged endings are present at both P0 and P3 (Figure 4K). These findings indicate that Netrin-G1 controls lanceolate complex formation during neonatal development.

We also tested whether the reduced number of lanceolate endings in *Ntng1*^*-/-*^ mice results from fewer axonal branches per neuron, a deficit in lanceolate complex formation in individual branches, or both. To address this, sparse neuronal labeling in *Ntng1*^*-/-*^ mice was performed to visualize the morphological features of individual Aβ RA-LTMRs and Aδ-LTMR neurons in hairy skin at P3 (Figures S3A). Interestingly, *Ntng1*^*-/-*^ mice exhibited a reduction in the number of innervated hair follicles per neuron (Figures S3B), consistent with the delay of guard hair wrapping by NFH^+^ endings observed at P0-P1 (Figures 4I). However, the number of branch points and area of individual neurons in *Ntng1*^*-/-*^ mice did not differ from controls (Figures S3C and S3D). Thus, while dispensable for initial stages of Aβ RA-LTMR and Aδ-LTMR skin innervation and axonal branching patterns, Netrin-G1 controls LTMR axonal wrapping around hair follicle and lanceolate complex formation during neonatal and early postnatal development.

The hairy skin lanceolate ending deficits in *Ntng1*^*-/-*^ mice prompted us to ask whether other LTMR end organs exhibit morphogenesis deficits in these mice (Handler and Ginty, 2021; Zimmerman et al., 2014). Interestingly, Meissner corpuscles in *Ntng1*^*-/-*^ mice appeared larger and their NFH^+^ axons displayed more enlargements than littermate controls (Figures 4L-4N), although the density of Meissner corpuscles was not altered in the mutants (Figure 4O). In wildtype Pacinian corpuscles, NFH^+^ Aβ RA-LTMR axons usually have one bulbous ultraterminal region (Handler and Ginty, 2021; Spencer and Schaumburg, 1973). However, Pacinian-innervating Aβ RA-LTMR axons in *Ntng1*^*-/-*^ mice exhibited an increase in the number of enlargements in their ultraterminal region (Figures 4L and 4P). These axonal enlargements resemble those observed in the hair follicle lanceolate endings, suggesting a common role for Netrin-G1 signaling in the morphogenesis of Aβ RA-LTMR axons and their end organs.

### Netrin-G1 functions in somatosensory neurons to regulate the formation of lanceolate endings and Aβ RA-LTMR-innervating corpuscles

To determine if Netrin-G1 functions in somatosensory neurons to promote lanceolate ending formation, Netrin-G1 was conditionally deleted from embryonic somatosensory neurons using *Advillin*^*Cre*^ mice (Hasegawa et al., 2007) and a conditional “ floxed” *Ntng1* allele (Zhang et al., 2016b). To increase the efficiency of embryonic *Ntng1* deletion, we generated *Advillin*^*Cre*^*;Ntng1*^*f/-*^ mutants and compared them to littermate controls carrying one copy of the null allele. As with *Ntng1*^*-/-*^ mice (Figure 4I), at P1, *Advillin*^*Cre*^*;Ntng1*^*f/-*^ conditional mutants showed a delay in wrapping of lanceolate endings around guard hairs (Figures 5A and 5B). Similarly, the conditional mutants displayed fewer NFH^+^ lanceolate endings and many enlarged endings associated with lanceolate complexes around guard hairs (Figures 5C and 5D). These findings suggest that Netrin-G1 acts in somatosensory neurons to promote lanceolate complex formation. Moreover, as in *Ntng1*^*-/-*^ mice, *Advillin*^*Cre*^*;Ntng1*^*f/f*^ mice also exhibited an increase in the area of Meissner corpuscles and the number of axonal enlargements (Figures 5E-5G) as well as an increase in the number of enlargements in axons in Pacinian corpuscles (Figures 5H and 5I). These findings indicate that Netrin-G1 acts in Aβ RA-LTMRs during formation of lanceolate endings associated with hair follicles, Meissner corpuscles of glabrous skin, and Pacinian corpuscles associated with the periosteum of bones.

**Figure 5.**
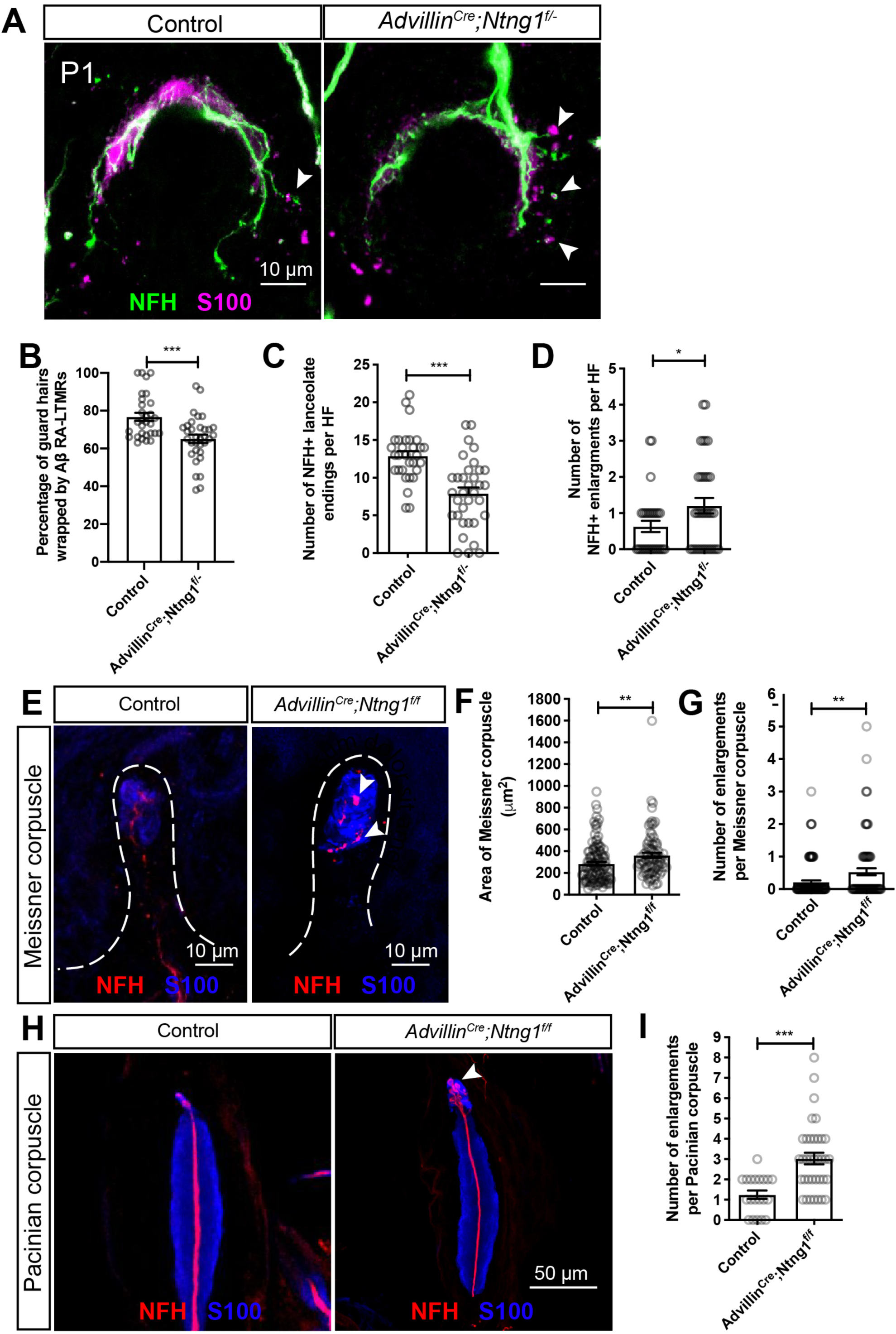
Netrin-G1 functions in somatosensory neurons to regulate mechanosensory end organ formation. **(A)** Whole-mount IHC images of guard hairs in back hairy skin of P1 control (*Ntng1*^*+/-*^) and *Advillin*^*Cre*^; *Ntng1*^*f/-*^ animals. **(B)** Quantification of the percentage of guard hairs wrapped by NFH^+^ lanceolate endings of P1 control and *Advillin*^*Cre*^; *Ntng1*^*f/-*^ animals. **(C and D)** Quantification of the number of NFH^+^ lanceolate endings (C) and the number of enlarged endings (D) per guard hair in P1 control (n = 3 animals) and *Advillin*^*Cre*^;*Ntng1*^*f/-*^ (n = 3 animals) mice. **(E)** Representative IHC images of forepaw glabrous skin sections, showing Meissner corpuscles in control and *Advillin*^*Cre*^; *Ntng1*^*f/f*^ mice. Arrowheads point to axonal enlargements. **(F and G)** Quantification of the area (F) and number of enlargements (G) of Meissner corpuscles in the epidermis of control (n = 3 animals) and *Advillin*^*Cre*^; *Ntng1*^*f/f*^ (n = 3 animals) mice. **(H and I)** Representative IHC images of Pacinian corpuscles (H) and quantification of the number of enlargements per Pacinian corpuscle (I) in control (n = 2 animals) and *Advillin*^*Cre*^;*Ntng1*^*f/f*^ (n = 3 animals) mice. The arrowhead points to axonal enlargements. Each dot represents a single Pacinian corpuscle. Student’s unpaired t test. *p<0.05, **p<0.01, ***p<0.001

### *Lrrc4c* is expressed in myelinating Schwann cells and terminal Schwann cells

Netrin-G ligand-1 (NGL-1), encoded by the gene *Lrrc4c*, is a type I transmembrane protein and a postsynaptic density (PSDs)-95-interacting postsynaptic adhesion molecule that interacts with Netrin-G1 (Lin et al., 2003; Choi et al., 2019; Matsukawa et al., 2014). NGL-1 selectively binds to Netrin-G1 to activate downstream signaling that promotes synapse formation (Kim et al., 2006; Seiradake et al., 2011). We therefore tested the hypothesis that NGL-1 functions as a ligand for Netrin-G1 in LTMR ending formation by determining the localization of NGL-1 and whether mice lacking NGL-1 exhibit the same LTMR deficits observed in *Ntng1* mutants. For these questions, we used *Lrrc4c* mutant mice in which the NGL-1 protein coding exon 3 of *Lrrc4c* was replaced with a β-geo cassette (Choi et al., 2019). This targeting strategy allowed us to examine the expression pattern of *Lrrc4c* in the skin by immunostaining for β-Gal. β-Gal signal was detected in S100^+^ TSCs and myelinating Schwann cells in hairy skin (Figure 6A). In complementary experiments, single-molecule RNA fluorescent *in situ* hybridization (smRNA-FISH) was performed on P3 hairy skin using probes against *Lrrc4c* and the glial specific gene *Plp1. Lrrc4c* signal was observed in close proximity to the *Plp1* signal around hair follicles and in the sensory nerve bundles near hair follicles (Figure 6B). These findings indicate that *Lrrc4c* is expressed in TSCs and myelinating Schwann cells when its receptor Netrin-G1 is expressed in LTMR axons, during the period of Netrin-G1-dependent end organ formation.

**Figure 6.**
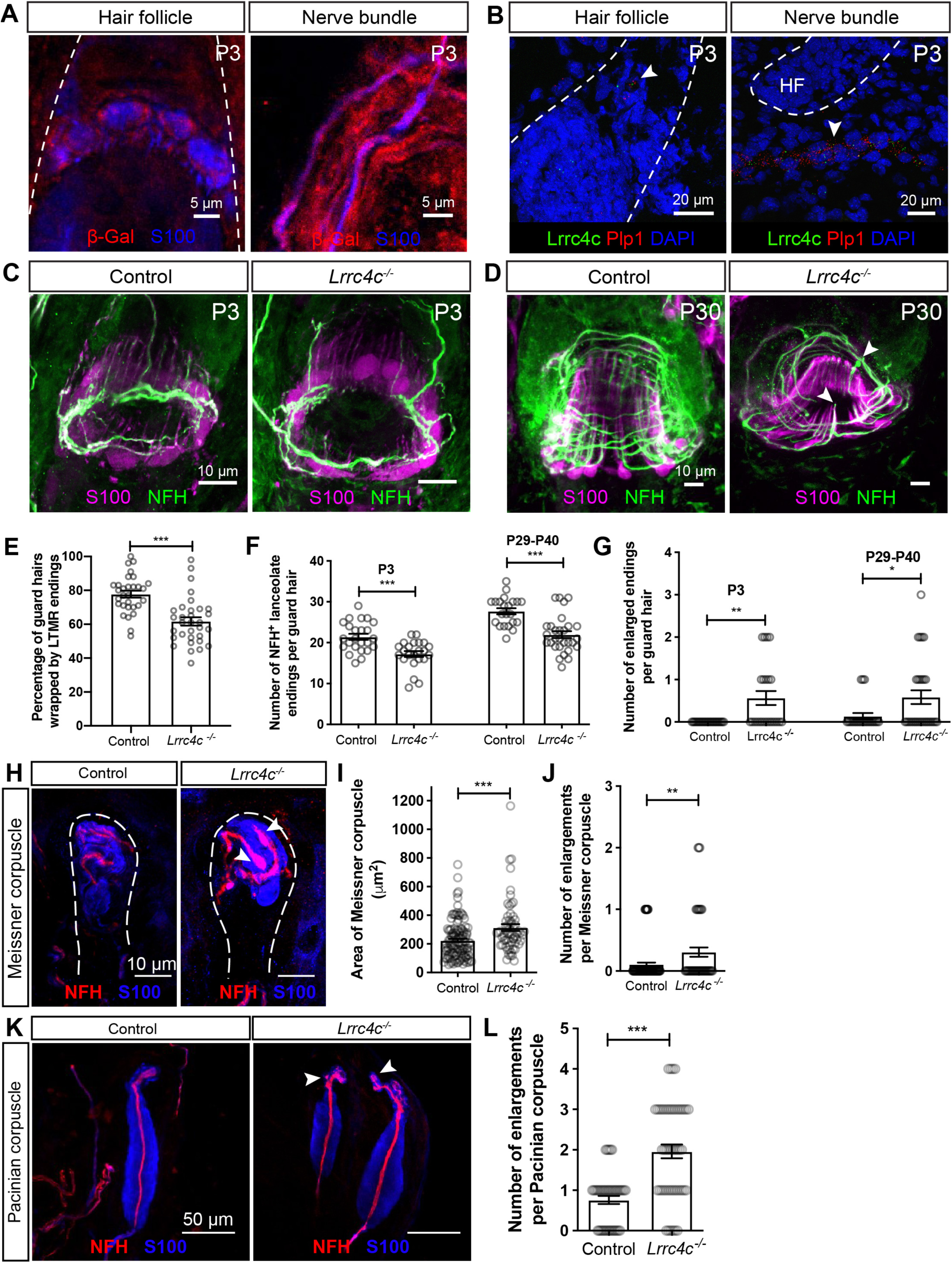
NGL-1 is expressed in terminal Schwann cells and contributes to the formation of lanceolate endings, Meissner corpuscles and Pacinian corpuscles. **(A)** Wholemount IHC images of P3 hairy skin sections, showing that β-Gal (red) is detected in S100^+^ terminal Schwann cells and non-myelinating Schwann cells (blue). Hair follicles are outlined with white dotted lines. Skin from wildtype animals was used as negative controls and no β-Gal signal was observed. **(B)** Example images of smRNA-FISH for P3 hairy skin sections. White arrowheads denote the detection of *Lrrc4c* mRNA (green) near *Plp1* mRNA (red). Hair follicles are outlined with white dotted lines. DAPI staining labels nuclei. **(C and D)** Whole-mount immunostaining images of guard hairs in back hairy skin of P3 (C) and P30 (D) control and *Lrrc4c*^*-/-*^ animals. **(E)** Quantification of the percentage of guard hairs wrapped by NFH^+^ lanceolate endings in P0 control and *Lrrc4c*^*-/-*^ animals (n = 3 animals per genotype). Each dot represents the result for one guard hair. **(F and G)** Quantification of the number of NFH^+^ lanceolate endings (F) and the number of enlarged endings (G) in control (n = 4 animals for P3; n = 3 animals for P29-P40) and *Lrrc4c*^*-/-*^ (n = 3 animals for P3; n = 4 animals for P29-P40) animals. **(H)** Representative IHC images of forepaw glabrous skin sections of control and *Lrrc4c*^*-/-*^ animals. **(I and J)** Quantification of the area (I) and number of enlargements (J) of Meissner corpuscles in the control and *Lrrc4c*^*-/-*^ mice (n = 4 animals for each genotype). **(K and L)** Representative IHC images (K) and quantification of the number of enlargements per Pacinian corpuscle (L) of adult control (n = 3 animals) and *Lrrc4c*^*-/-*^ mice (n = 3 animals). Each dot represents a single Pacinian corpuscle. Student’s unpaired t test. *p<0.05, **p<0.01, ***p<0.001

### *Lrrc4c* regulates LTMR ending formation

To determine if *Lrrc4c* also regulates formation of lanceolate complexes, Meissner corpuscles, and Pacinian corpuscles, we examined hairy and glabrous skin of control and *Lrrc4c*^*-/-*^ mice at different ages (Figures 6C-6J). Similar to *Ntng1*^*-/-*^ mice, *Lrrc4c*^*-/-*^ mice showed a delay in the wrapping of lanceolate endings around guard hairs at P0 (Figure 6E). In both P3 pups and young adults (P29-P40), lanceolate complexes around guard hairs in *Lrrc4c*^*-/-*^ mice had fewer lanceolate endings, compared to those of littermate controls (Figures 6C, 6D and 6F). The number of enlarged lanceolate endings was also increased in *Lrrc4c*^*-/-*^ mice (Figures 6G). We note that the lanceolate ending phenotype is milder than that observed in *Ntng1*^*-/-*^ mice, suggesting that additional ligands might signal through Netrin-G1 to promote lanceolate ending formation. Furthermore, sparse labeling of individual Aβ RA-LTMR and Aδ-LTMR neurons in P3 hairy skin revealed that *Lrrc4c*^*-/-*^ mice phenocopied the hair follicle innervation delay observed in *Ntng1*^*-/-*^ mice (Figures S3E-S3H). In addition, *Lrrc4c*^*-/-*^ mice exhibited an increase in the area and number of enlargements of Meissner corpuscles (Figures 6H-6J) as well as an increase in the number of enlargements in Pacinian corpuscles (Figures 6K and 6L), although these phenotypes were also milder that those observed *Ntng1*^*-/-*^ mice. These findings together with the *Lrrc4c* expression pattern suggest that NGL-1 in TSCs signals through Netrin-G1 in LTMR axons to promote formation of lanceolate ending complexes as well as Meissner corpuscles and Pacinian corpuscles.

## Discussion

Hair follicle lanceolate complexes are mechanically sensitive end organs that transform hair deflection into LTMR electrical signals. Lanceolate complexes are formed by Aβ RA-LTMR, Aδ-LTMR, and C-LTMR axonal endings and their associated TSCs. How this unique complex forms during development and the molecular players involved have been unclear. In this study, we combined genetic labeling of LTMRs and TSCs, histological analyses, and genetic manipulations to characterize developmental steps leading to the formation of Aβ RA-LTMR and Aδ-LTMR lanceolate ending complexes. We found that Aβ RA-LTMRs innervate hair follicles earlier than Aδ-LTMRs, and that branches of both LTMR subtypes exhibit robust pruning. Nascent lanceolate endings are associated closely with TSCs during lanceolate complex formation. Moreover, Netrin-G1 in sensory neurons acts to promote the formation of lanceolate complexes around hair follicles as well as Meissner corpuscles and Pacinian corpuscles innervated by Aβ RA-LTMRs. *Lrrc4c*, encoding NGL-1, a ligand for Netrin-G1, is expressed in TSCs and myelinating Schwann cells, and loss of *Lrrc4c* led to similar albeit less dramatic deficits in LTMR endings, revealing a critical role for neuron-glial interactions and NGL-1– Netrin-G1 signaling in LTMR end organ formation.

### Pruning of touch sensory axons during postnatal development

Across the nervous system, functional circuits are established through a progressive series of developmental processes, often including axonal branch elaboration, and subsequently refined by regressive processes such as the pruning of excess axonal branches (Furusawa and Emoto, 2021). Developmental neurite and synaptic pruning have been observed in many neuronal subtypes (Luo and O’Leary, 2005; Riccomagno and Kolodkin, 2015); this ensures that unnecessary neurites or connections are removed to refine neural circuits for proper function. Using genetic labeling of developing Aβ RA-LTMRs and Aδ-LTMRs, we observed exuberant branching followed by pruning of touch mechanosensory neuron endings at two stages. First, excess axonal branches that do not innervate hair follicles are eliminated, which may contribute to the formation of precise functional receptive fields of LTMRs. Second, over-produced lanceolate endings that extend towards the basal side of the skin are pruned, which may ensure high sensitivity, and directional selectivity in the case of Aδ-LTMRs (Rutlin et al., 2014), to hair deflections (Handler and Ginty, 2021). The common pruning strategies across these two LTMR types suggest a common mechanism instructing this regressive event. If this is the case, then C-LTMRs, which also form lanceolate endings in hairy skin, may exhibit similar types of pruning during early postnatal development, and future experiments targeting developing C-LTMRs may help address this question. It is possible that local neurotrophic factors in the skin, such as keratinocyte-derived BDNF (Rutlin et al., 2014), instruct the survival and maintenance of axons that innervate hair follicles. Another intriguing question is whether sensory neuron activity plays a role in instructing pruning. The axonal pruning observed takes place around and shortly after birth, which roughly coincides with the period of time when evoked sensory activity in the mechanosensory system begins (Meltzer et al., 2021). It will therefore be interesting to determine if mechanosensory neuron activity contributes to pruning of LTMR endings in hairy skin.

### The role of Netrin-G1 signaling in lanceolate complex formation

The close association between developing TSCs and LTMRs suggests that TSCs may be instrumental in promoting LTMR axonal branching and lanceolate complex formation. Indeed, Netrin-G1 receptors and at least one Netrin-G1 ligand, TSC-derived NGL-1, act to promote lanceolate ending formation. Interestingly, a recent finding shows that microglia accumulation around subcerebral projection axons depends on NGL-1–Netrin-G1 signaling (Fujita et al., 2020), suggesting a general role for Netrin-G1 signaling in mediating neuron-glial cell communication. We speculate that, in addition to NGL-1, other ligands also signal through Netrin-G1, as *Lrrc4c* mutants exhibited a weaker phenotype than the *Ntng1* mutants. How Netrin-G1 signaling could mediate extension and structural integrity of lanceolate endings remain to be determined. Because Netrin-G1 is a GPI-anchored protein without an intracellular domain (Nakashiba et al., 2000), it likely functions together with a co-receptor to activate downstream signaling within lanceolate processes. Such a putative Netrin-G1 co-receptor is likely expressed in LTMRs and thus present in our LTMR sequencing datasets (Sharma et al., 2020; Zheng et al., 2019). Nevertheless, the discovery of an essential role of Netrin-G1 signaling in lanceolate endings, Meissner corpuscles, and Pacinian corpuscles establishes a critical role for neuron-glia interactions and an inter-cellular molecular dialogue that instruct the development of cutaneous mechanosensory end organs.

## Acknowledgements

We thank all the Ginty lab members for helpful discussions and feedback on this manuscript. We thank the Harvard Medical School Neurobiology Imaging Facility for assistance with small-RNA FISH experiments. This research was supported by a Hanna Gray Fellowship (S.M.), a Stuart H.Q. and Victoria Quan fellowship (Q.Z.), a Howard Hughes Medical Institute–Jane Coffin Childs Fellowship (A.H.), the Edward R. and Anne G. Lefler Center for Neurodegenerative Disorders (D.D.G.), and NIH R35 NS097344 (D.D.G.). D.D.G. is an investigator of the Howard Hughes Medical Institute. This article is subject to HHMI’s Open Access to Publications policy. HHMI lab heads have previously granted a nonexclusive CC BY 4.0 license to the public and a sublicensable license to HHMI in their research articles. Pursuant to those licenses, the author-accepted manuscript of this article can be made freely available under a CC BY 4.0 license immediately upon publication.

## Author contributions

S.M. and D.D.G. designed the study; S.M., K.C, E.A., A.H., Q.Z., and C.S. performed experiments; C.S. and S.I. provided tissues and mice for experiments. S.M., K.C, E.A., A.H., Q.Z., and D.D.G. analyzed data; and S.M., K.C, E.A., A.H., Q.Z., C.S., S.I. and D.D.G. wrote the paper.

---

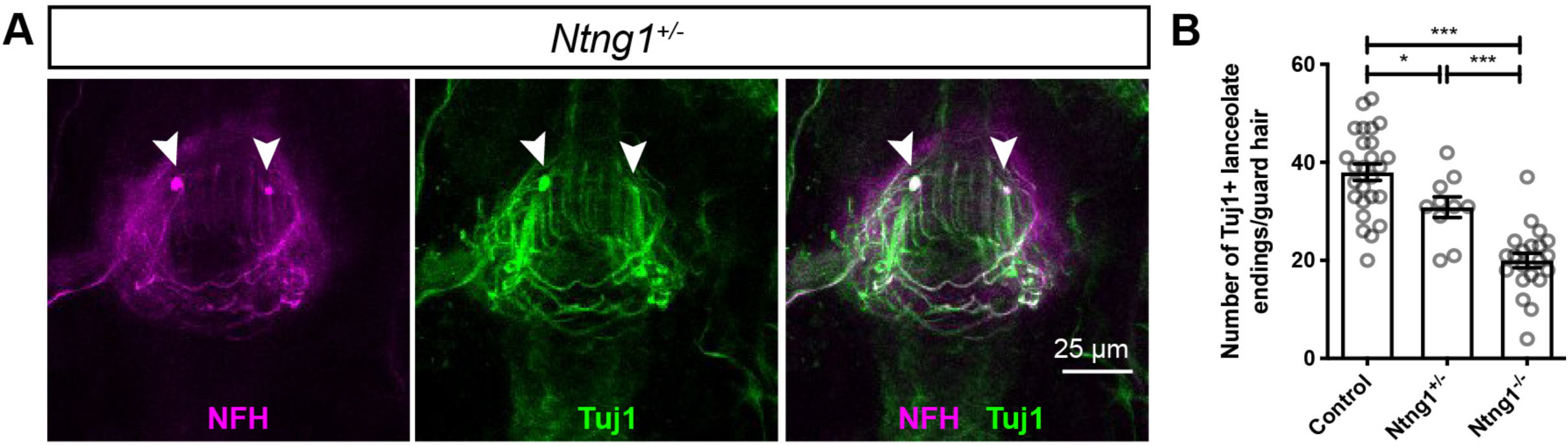

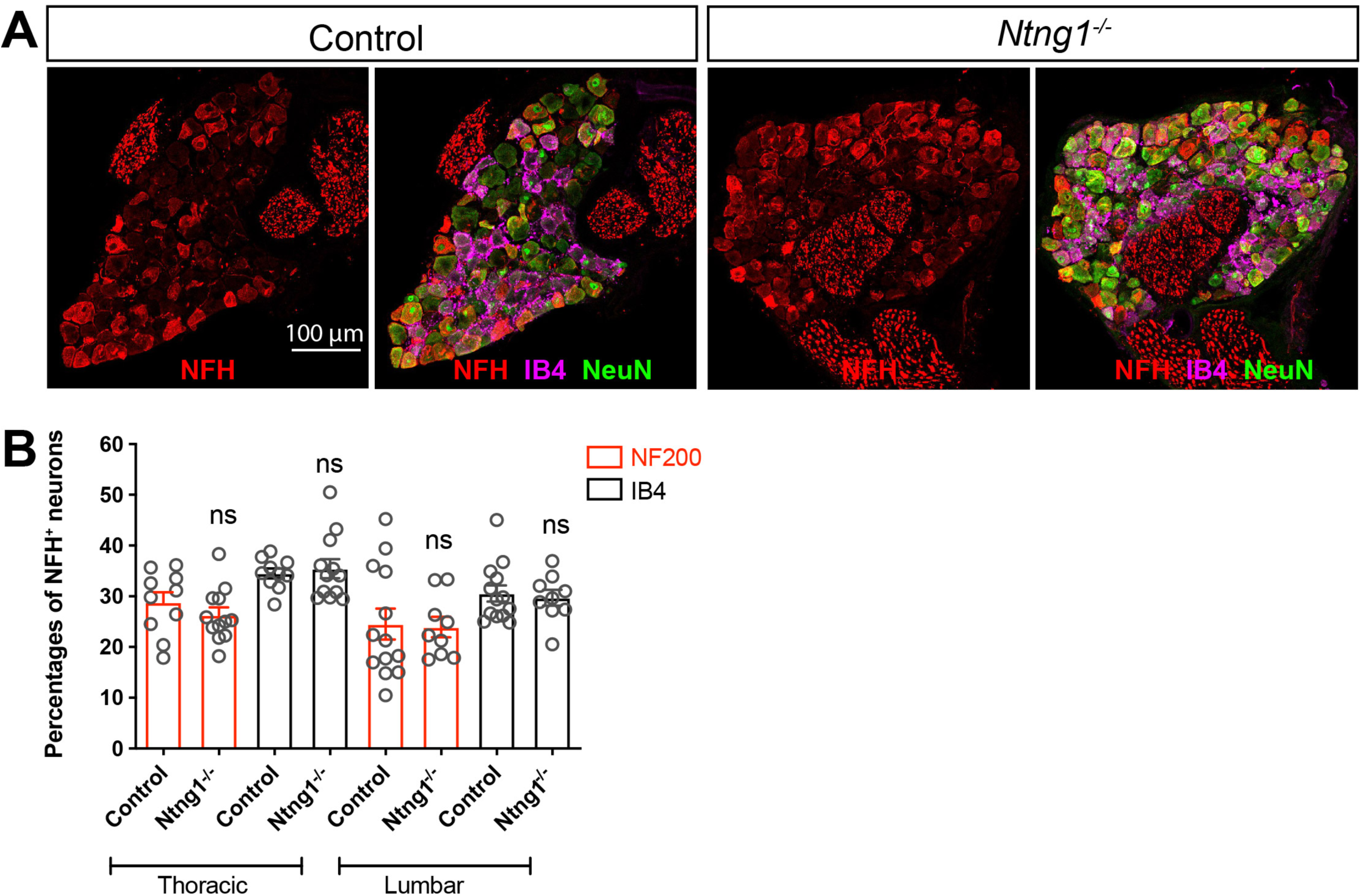

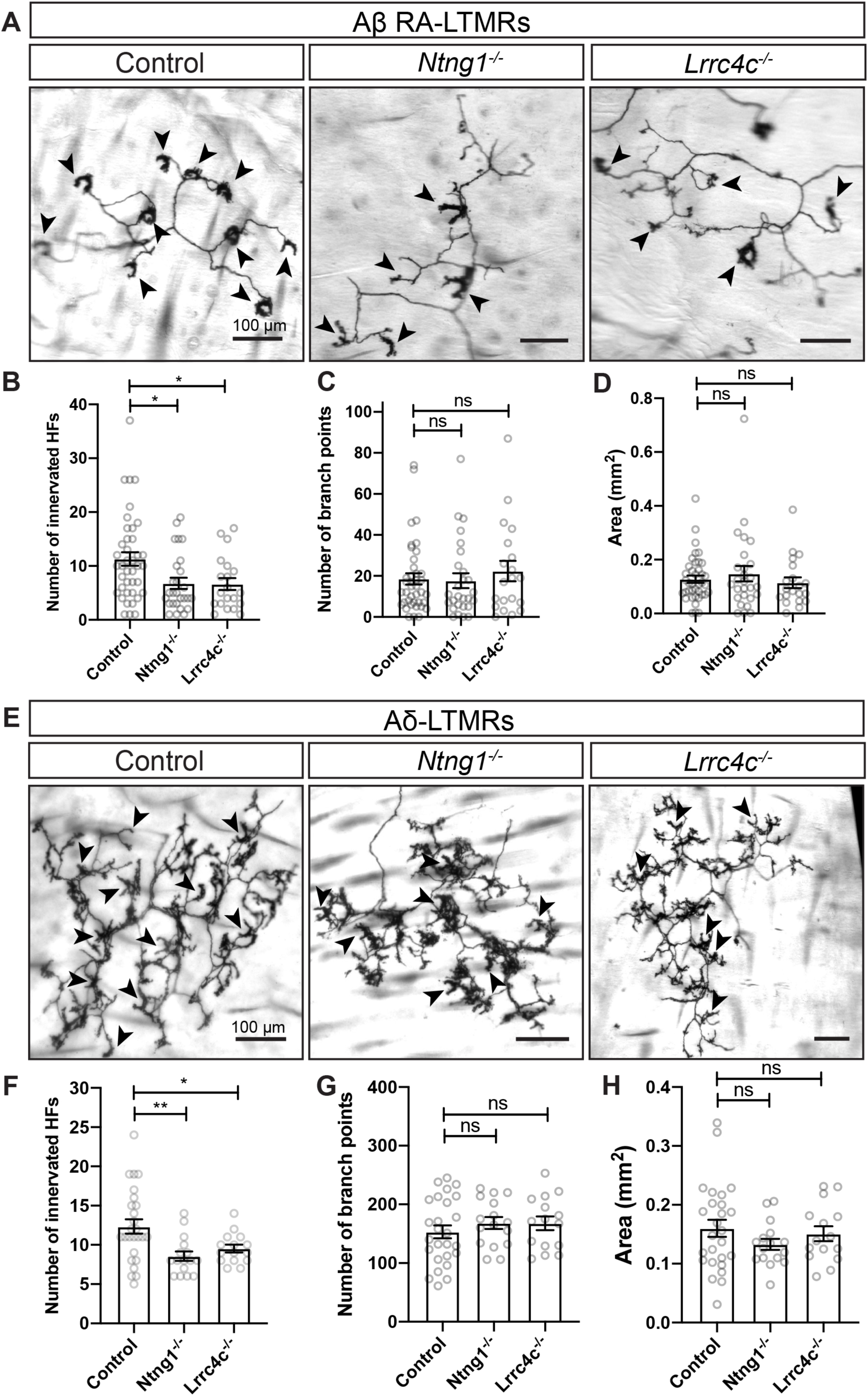

## Notes

### Competing Interest Statement

The authors have declared no competing interest.

