## Supplemental text for "A role for axon-glial interactions and Netrin-G1 signaling in the formation of low-threshold mechanoreceptor end organs"

**Supplementary Information Text**

**Experimental Procedures**

Mouse strains

All procedures were conducted according to animal protocols approved by the Harvard Medical School Institutional Animal Care and Use Committee (IACUC). Mouse lines used include *Advillin^Cre^* (Hasegawa et al., 2007); *Ret^CreER^* (Luo et al., 2009); *TrkB^CreER^* (Rutlin et al., 2014); *R26^LSL-tdTomato^* (Ai14) (Jax #007908); *Brn3a^f(AP)^* (Badea et al., 2009); *R26^IAP^* (Jax Cat# 009253); *Netrin-G1^-/-^* (Nishimura-Akiyoshi et al., 2007); *Netrin-G1^flox^* (Zhang et al., 2016); *Lrrc4c^-/-^* (MMRRC #032444) (Choi et al., 2019). For histological experiments E15.5-P80 male and female mice were used. At least three animals per genotype were examined for each experiment, unless noted in the figure legends. In all experiments where mutants were compared to control mice, the experimenter was blind to genotype during data analyses.

Tamoxifen treatment

Tamoxifen was dissolved in ethanol (20 mg/ml), mixed with an equal volume of sunflower seed oil (Sigma), vortexed vigorously for 15 min and centrifuged under vacuum for 40 min to remove ethanol. The working solution (20mg/ml) was kept at -20°C and delivered to pregnant females via oral gavage. The morning after coitus was designated as E0.5 and the day of birth as P0.

Tissue collection and fixation

Animals P10 and older were anesthetized using isoflurane inhalation and a transcardial perfusion was performed using approximately 10-20 mL modified Ames Media with heparin, followed by 10-20 mL 4% paraformaldehyde (PFA) in 1xPBS. Animals younger than P6 were anesthetized in ice and sacrificed quickly by decapitation. Hair was removed from back hairy skin by application of a commercial hair remover. Spinal cords were post-fixed in 4% PFA overnight at 4°C, while legs, paws, and back hairy skin were post-fixed in Zamboni’s fixative (Fisher Cat# NC9335034) overnight at 4°C. The following morning, all samples were washed with 1xPBS for 3-6 hours approximately six times. All tissues were kept in 1xPBS with 0.01% sodium azide at 4°C for long-term storage.

Immunohistochemistry via cryo-sectioning

Tissues were dissected and cryoprotected in 30% sucrose at 4 °C for 1-2 nights before freezing with OCT (Fisher Cat# 1437365) over a dry ice and 100% ethanol bath. Samples were placed in labelled Eppendorf tubes and transferred to a -80°C freezer for at least an hour before sectioning. Spinal cord and DRG samples were sectioned at a thickness of 12-20 μm, while skin tissues were sectioned at 25-30 μm. The spinal cord and DRG sections on slides were left to dry at room temperature overnight and the skin tissue slides were left to dry at room temperature over 2-3 nights. Slides were transferred to a -20°C freezer.

For immunohistochemistry, slides were removed from the -20 °C freezer and allowed to warm up on the bench for 30 minutes. A hydrophobic barrier was drawn around the samples using a PAP pen and the slides were washed 3 times for 5 minutes each with 1xPBS. The tissue was then blocked in 0.1% Triton X-100 in 1xPBS (0.1% PBST) with 5% normal donkey or goat serum for an hour at room temperature, incubated with primary antibodies diluted in filtered 1xPBS with 5% serum overnight at 4°C, and washed 4 times for 5 minutes each with 0.02% Tween-20 in 1XPBS. Tissues were then incubated with secondary antibodies diluted in filtered 1xPBS with 5% serum for 2 hours at room temperature and washed 4 times for 5 minutes each with 0.02% Tween-20 detergent. Slides with hairy or glabrous skin tissue were mounted and coverslipped with DAPI Fluoromount-G (Fisher Cat# 0100-20), while spinal cord and DRG slides were mounted with Fluoromount Aqueous Mounting Medium (Sigma Cat# F4680). Slides were stored in a dark slide box at 4°C for at least overnight and up to a week before imaging.

The following primary antibodies and lectins and their dilutions were used for IHC in these experiments: goat anti-Netrin-G1 (R and D Systems Cat# AF1166, RRID:AB_2154822), Isolectin B4 647 (Invitrogen, RRID: SCR_014365, 1:500), chicken polyclonal anti-beta Galactosidase (Abcam, Cat# ab936, 1:500), chicken polyclonal anti-NFH (Aves, Cat# NFH0211, 1:500), rabbit anti-Tuj1 (BioLegend Cat# 802001, RRID:AB_2564645), mouse monoclonal anti-NeuN (Millipore, RRID: AB_2298772, 1:500), rabbit polyclonal anti-S100 Beta (Fisher Cat# 15146-1-AP, 1:500), and rat polyclonal anti-Troma1 (DSHB, RRID: AB_531826, 1:50). Different Invitrogen Alexa Fluor conjugated secondary antibodies were used at 1:500 dilution for these experiments in channels 405, 488, 546, and 647 and in donkey and goat host animals (Thermo Fisher Scientific).

Wholemount Immunostaining

Back hairy skin samples were cut into approximately 1 cm² pieces with a razor blade. The skin samples were washed every 30-60 minutes with 0.3% Triton X-100 in PBS (0.3% PBST) for 5-8 hours at room temperature, incubated with primary antibodies in 0.3% PBST with 5% serum and 20% DMSO for 3-5 days at room temperature. Skin tissues were then washed again every 30 minutes with 0.3% Triton X-100 (0.3% PBST) for 5-8 hours, incubated with secondary antibodies in 0.3% PBST with 5% serum and 20% DMSO for 2-3 days, and washed every 30-60 minutes with 0.3% PBST for 5-8 hours. Finally, the samples were dehydrated in 50%, 75%, and 100% methanol for 10 minutes each before transferring them to 100% methanol at 4°C overnight. Samples were stored in 100% methanol at -20°C for long-term storage before imaging. Skins were cleared with BABB (1 part benzyl alcohol: 2 parts benzyl benzoate) for 5 min. The skin tissue was then mounted on a glass slide and a coverslip for confocal imaging.

For Pacinian corpuscle staining, the interosseous membrane around the fibula was dissected from legs that were previously fixed overnight with Zamboni's fixative. These tissue samples were subjected to the same wholemount immunostaining protocol above except 1% PBST was used in place of 0.3% PBST to increase the antibody penetrance. Pacinian corpuscle data analysis was conducted using FIJI ImageJ software. The number of NFH-positive axonal enlargements and extraneous branches were quantified using the counting tool.

Whole-mount alkaline phosphatase staining

For single sensory axon analyses, *Brn3a^f(AP)^* or *R26^IAP^* mice were used to express alkaline phosphatase in genetically labeled sensory neurons. Aβ RA-LTMRs were labeled using *Ret^CreER^* treated with 0.1 - 0.5 mg tamoxifen at E11.5, and Aδ-LTMRs were labeled using *TrkB^CreER^* treated with 0.05 - 0.2 mg tamoxifen at E12.5-14.5. Whole-mount placental alkaline phosphatase staining was performed as previously described (Kuehn et al., 2019). Mice (E15.5, E17.5, P0, P3, P5, P10, and P21) were euthanized and hair was removed. Back hairy skins were dissected and post-fixed in Zamboni’s fixative at 4 °C overnight. Skins were then washed with PBS 3 times for 30 minutes at room temperature and placed in a 65 °C chamber for 2 hours. Skins were then washed with B3 buffer (0.1 M Tris pH 9.5, 0.1 M NaCl, 50 mM MgCl2) three times for 15 minutes at room temperature, and incubated in BCIP (Sigma Cat#11383221001)/NBT(Sigma Cat# 11383213001) buffer (3.4 μl/ml each) in B3 buffer with 0.1% Tween-20 for ~24 hours at room temperature. Tissues were then washed with PBS, fixed with 4% PFA/PBS for 1 hour at room temperature, rinsed with PBS, and dehydrated sequentially with 50%, 75%, and 100% ethanol overnight and cleared in BABB for imaging.

Meissner corpuscle analysis

Fingertips were dissected from the forepaws of each animal, cryo-protected, frozen, and sectioned at 25 μm thickness. Tissues were stained for Neurofilament (NFH), S100b, and DAPI using the method above, and imaged with confocal microscopy. Meissner corpuscle data analysis was conducted on ImageJ software. Briefly, individual Meissner corpuscles were identified by the presence of S100b signal in the dermis in a corpuscular shape. To assess the size of individual Meissner corpuscles, two measurements were taken for each individual corpuscle: width and area. To assess the width of a corpuscle, the ImageJ straight-line tool was used to draw a line across the width of the corpuscle parallel to the surface of the skin and the ImageJ ‘Measure’ feature was used to obtain the width. Similarly, to assess the area, the polygon tool was used to draw a boundary around the S100b^+^ corpuscle and the “Measure” plugin was used to obtain the area measurement. The data was compiled and compared across genotypes and ages.

Tissue processing for RNA-FISH

For sample preparation, back hairy skins from mice were rapidly dissected and frozen in dry ice-cooled 2-metylbutane and stored at −80 °C. Skins were sectioned at −20 °C at a thickness of 15 μm and RNAs were detected by RNAscope (Advanced Cell Diagnostics) using the manufacturer’s protocol. Nuclei were stained with DAPI. Mm-Plp1 (Cat#: 428188-C2) and Mm-Lrrc4c (Cat# 490331) probes were used.

Electron microscopy

Electron microscopy of the back hairy skin was conducted as previously described (Neubarth et al., 2020; Zhang et al., 2019). Briefly, guard hairs on the back hairy skin were identified based on their morphological appearance and isolated by trimming all surrounding hairs. Skin tissues were excised and immersed in 2% PFA (Electron Microscopy Sciences) and 2.5% glutaraldehyde (Electron Microscopy Sciences) for 1 hour at room temperature, further dissected, and subsequently fixed in the same fixative overnight at 4 °C. These tissues were then osmicated with reduced osmium (1% osmium tetroxide (Electron Microscopy Sciences)/1.5% potassium ferrocyanide (MilliporeSigma)), stained with 1% uranyl acetate (Electron Microscopy Sciences), dehydrated with an ethanol series followed by propylene oxide, and infiltrated and embedded with an epoxy resin mix (LX-112, Ladd Research). Samples were then cured in the oven at 60 °C for 48-96 hours.

Samples were sectioned using a Leica EM UC7 ultramicrotome with Diatome diamond knives. Guard hairs were identified in semi-thin sections (1 μm), and ultrathin sections (40-60 nm) of the same regions were picked up on glow discharged formvar/carbon films on slot grids (Ted Pella). Ultrathin sections were imaged using a JEOL 1200EX transmission electron microscope at 80 kV accelerating voltage.

Statistical methods

All data are expressed as the mean +/- standard error of the mean (SEM). The number of animals per group used in each experiment is denoted in the figure legend. Comparisons between two groups in all experiments were performed using Student’s t test. One-way ANOVA was used in the case of three or more groups of the same condition were compared. Two-way ANOVA was used when at least two groups with multiple conditions were compared. For ANOVA tests, post hoc comparisons were performed using the post hoc test indicated in the figure legend. The p values of post hoc comparisons are denoted with asterisks in the figures. Unless otherwise noted, *, p < 0.05, **, p<0.01 and ***, p<0.001. All statistics were performed using the GraphPad Prism software.

**Supplemental Figure Legends**

**Figure S1. Netrin-G1 heterozygous mutants show an intermediate phenotype**

**(A)** Whole-mount immunostaining images of guard hairs in back hairy skin in *Ntng1^+/-^* animals. Aβ RA-LTMRs lanceolate endings are marked by NFH (magenta) and Tuj1 (green) labeling.

**(B)** Quantification of the number of Tuj1^+^ lanceolate endings in control (n = 7 animals), *Ntng1^+/-^* (n = 2 animals) and *Ntng1^-/-^* (n = 3 animals) animals. Each dot represents a single hair follicle. One-way ANOVA test. *p<0.05, ***p<0.001.

**Figure S2. No cell loss was observed in Netrin-G1 mutant mice**

**(A)** IHC images of DRG sections from control and *Ntng1^-/-^* animals (n = 3 animals per genotype) at P40. Large and medium diameter neurons are labeled with NFH. IB4 labels non-peptidergic neurons and NeuN labels all neurons.

**(B)** No significant differences were observed between the percentages of DRG neurons that are NFH^+^ or IB4^+^ in control and *Ntng1^-/-^* animals. Each dot represents a DRG section. Student’s unpaired t test. ns, not significant.

**Figure S3. NGL-1**–**Netrin-G1 signaling regulates hair follicle innervation by Aβ RA-LTMRs and Aδ-LTMRs**

**(A)** Whole-mount alkaline phosphatase staining images of hairy skin reveal peripheral terminals from individual Aβ RA-LTMRs in control, *Ntng1^-/-^* and *Lrrc4c^-/-^* animals at P3. Arrowheads point to axons that innervate hair follicles.

**(B-D)** Quantification of the number of hair follicles innervated (B), the number of branch points (C) and the area (D) of individual Aβ RA-LTMRs in control (n = 3 animals), *Ntng1^-/-^* (n = 2 animals) and *Lrrc4c^-/-^* (n = 2 animals) animals. Each dot represents a single neuron. One-way ANOVA test.

**(E)** Whole-mount alkaline phosphatase staining images of individual hairy skin Aδ-LTMRs in control, *Ntng1^-/-^* and *Lrrc4^-/-^* animals at P3. Arrowheads point to axons that innervate hair follicles.

**(F-H)** Quantification of the number of hair follicles innervated (F), the number of branch points (G) and the area (H) of individual Aδ-LTMRs in control, *Ntng1^-/-^* and *Lrrc4c^-/-^* animals (n = 3 animals per genotype). Each dot represents a single neuron. One-way ANOVA test.

ns, not significant; *p < 0.05; **p < 0.01.
